# One thousand soils for molecular understanding of belowground carbon cycling

**DOI:** 10.1101/2022.12.12.520098

**Authors:** Maggie M. Bowman, Alexis E. Heath, Tamas Varga, Anil K. Battu, Rosalie K. Chu, Jason Toyoda, Tanya E. Cheeke, Stephanie S. Porter, Kevan Moffett, Brittany Letendre, Odeta Qafoku, John R. Bargar, Douglas Mans, Nancy Hess, Emily B. Graham

**Affiliations:** Pacific Northwest National Laboratory, Richland, WA, 99354, USA; Bowling Green State University, Bowling Green, OH, 43403, USA; School of Biological Sciences, Washington State University, Richland, WA, USA; School of the Environment, Washington State University, Vancouver, WA 98686, USA

## Abstract

While significant progress has been made in understanding global carbon (C) cycling, the mechanisms regulating belowground C fluxes and storage are still uncertain. New molecular technologies have the power to elucidate these processes, yet we have no widespread standardized implementation of molecular techniques. To address this gap, we introduce the Molecular Observation Network (MONet), a decadal vision from the Environmental Molecular Sciences Laboratory (EMSL), to develop a national network for understanding the molecular composition, physical structure, and hydraulic and biological properties of soil and water. These data are essential for advancing the next generation of multiscale Earth systems models. In this paper, we discuss the 1000 Soils Pilot for MONet, including a description of standardized sampling materials and protocols and a use case to highlight the utility of molecular-level and microstructural measurements for assessing the impacts of wildfire on soil. While the 1000 Soils Pilot generated a plethora of data, we focus on assessments of soil organic matter (SOM) chemistry via Fourier-transform ion cyclotron resonance-mass spectrometry and microstructural properties via X-ray Computed Tomography to highlight the effects of recent fire history in forested ecosystems on belowground C cycling. We observed decreases in soil respiration, microbial biomass, and potential enzyme activity in soils with high frequency burns. Additionally, the nominal oxidation state of carbon in SOM increased with burn frequency in surface soils. This results in a quantifiable shift in the molecular signature of SOM and shows that wildfire may result in oxidation of SOM and structural changes to soil pore networks that persist into deeper soils.

## Introduction

Soil organic matter (SOM) is a critical part of the global carbon (C) cycle. Belowground ecosystems contain more C than stored in terrestrial vegetation and the atmosphere combined (Crowther et al., 2019; Crowther et al., 2016; Jobbágy and Jackson, 2000), and SOM is the largest and most biologically active portion of soil C. SOM decomposition is regulated by a complex and interacting set of factors including soil structure, moisture distribution, temperature, pH, and nutrient status; collectively, these factors determine accessibility, bioavailability, and rate kinetics of SOM (Graham and Hofmockel, 2022). Despite the importance of SOM in the global C cycle, the drivers of SOM decomposition from molecular to continental scales are not well understood.

Recent research suggests that standardized, spatially-resolved, and high-resolution data may be critical to reducing the uncertainty surrounding estimates of belowground C dynamics under future climate scenarios (Lehmann and Kleber, 2015; Viscarra Rossel et al., 2019). For example, pore-scale structural characterization could reveal spatial factors that inhibit SOM decomposition, and molecular chemistry may provide insight into SOM reactivity that cannot be observed from bulk measurements. As of yet, such information is lacking from soil databases (e.g., Web Soil Survey, International Soil Radiocarbon Database, and Soils Data Harmonization), and no widespread implementations of standardized high-resolution techniques simultaneously investigate the myriad factors influencing SOM decomposition (Bond-Lamberty and Thomson, 2010; Lawrence et al., 2020; NRCS; Wieder et al., 2021).

Consequently, soil C cycles are poorly represented in biogeochemical and Earth System Models (ESMs) that predict changes in the global climate (Bradford et al., 2016). For example, most ESMs simulate changes in belowground C concentration within coarsely defined pools (i.e., agnostic of SOM composition) rather than representing chemical and microbial reactions within SOM pools (Bond-Lamberty et al., 2020; Malhotra et al., 2019; Rasmussen et al., 2018; Waring et al., 2020). The depth of soil molecular and microphysical/chemical information presents a major limitation to improving models at regional and Continental United States (CONUS) scales, where impacts of climate change are manifested and can be understood.

Led by a team at the Environmental Molecular Science Laboratory (EMSL), a new decadal initiative is developing a CONUS-scale database of soil compositional, physical, and metagenomic measurements that are open and FAIR (Findable, Accessible, Interoperable and Reusable), termed the Molecular Observation Network (MONet) program (Todd-Brown et al., 2022; Wilkinson et al., 2016). The goal of MONet is to facilitate molecular data syntheses and modelling activities that can reduce uncertainty in soil C simulation at regional and CONUS scales. In this article, we describe the 1000 Soils Pilot for MONet, which focuses on understanding the drivers and fluxes of soil C cycles through standardized molecular measurements. It represents our initial goal to collect soil cores from 1000 locations.

To highlight the power of high-resolution measurements for understanding soil structure, molecular SOM composition, and SOM decomposition, we also present a use case from the 1000 Soils Pilot. Because wildfires are becoming increasingly prevalent worldwide (Abatzoglou and Williams, 2016; Aponte et al., 2016) and have lasting impacts on ecosystem structure and function (Abatzoglou and Williams, 2016; Certini, 2005; Doerr et al., 2006; Lombao et al., 2021; Matosziuk et al., 2020; Neary et al., 2005; Verma and Jayakumar, 2012; Wang et al., 2012), we focus our use case on a set of three locations with varied burned histories. We show that wildfire may result in oxidation of SOM and in structural changes to soil pore networks that persist into deeper soils, which are not investigated by typical sampling schema. These are important considerations in the management of forest ecosystem restoration post-wildfire and present an interesting avenue for more robust investigations as the MONet database expands.

### Approach

Through partnerships with individual researchers and ecological networks (Figure 1A), the 1000 Soils Pilot facilitated the collection of 76 sets of soil cores using a standardized sampling kit and field protocols. Collaborators collected two 30-cm (3-inch dia.) replicate intact cores using a slide hammer soil corer (AMS, Inc., USA), as well as four 10-cm (2-inch dia.) soil cores using a wooden block and mallet. Other field-based measurements included soil temperature, moisture, and electrical conductivity, measured using a Teros 12 Probe and ZSC Bluetooth interface (Meter Group, Inc., USA). Soil infiltration rate was measured using the single-ring infiltrometer method (Dane and Topp, 2020; Klute and Dirksen, 1986). A step-by-step sampling protocol is available in the SI and at protocols.io. After sample collection, all data was generated by EMSL or EMSL-contracted vendors. All data from the 1000 Soils Pilot are available via Zenodo (https://zenodo.org/communities/emsl-monet; Bowman et al., 2022).

**Figure 1:**
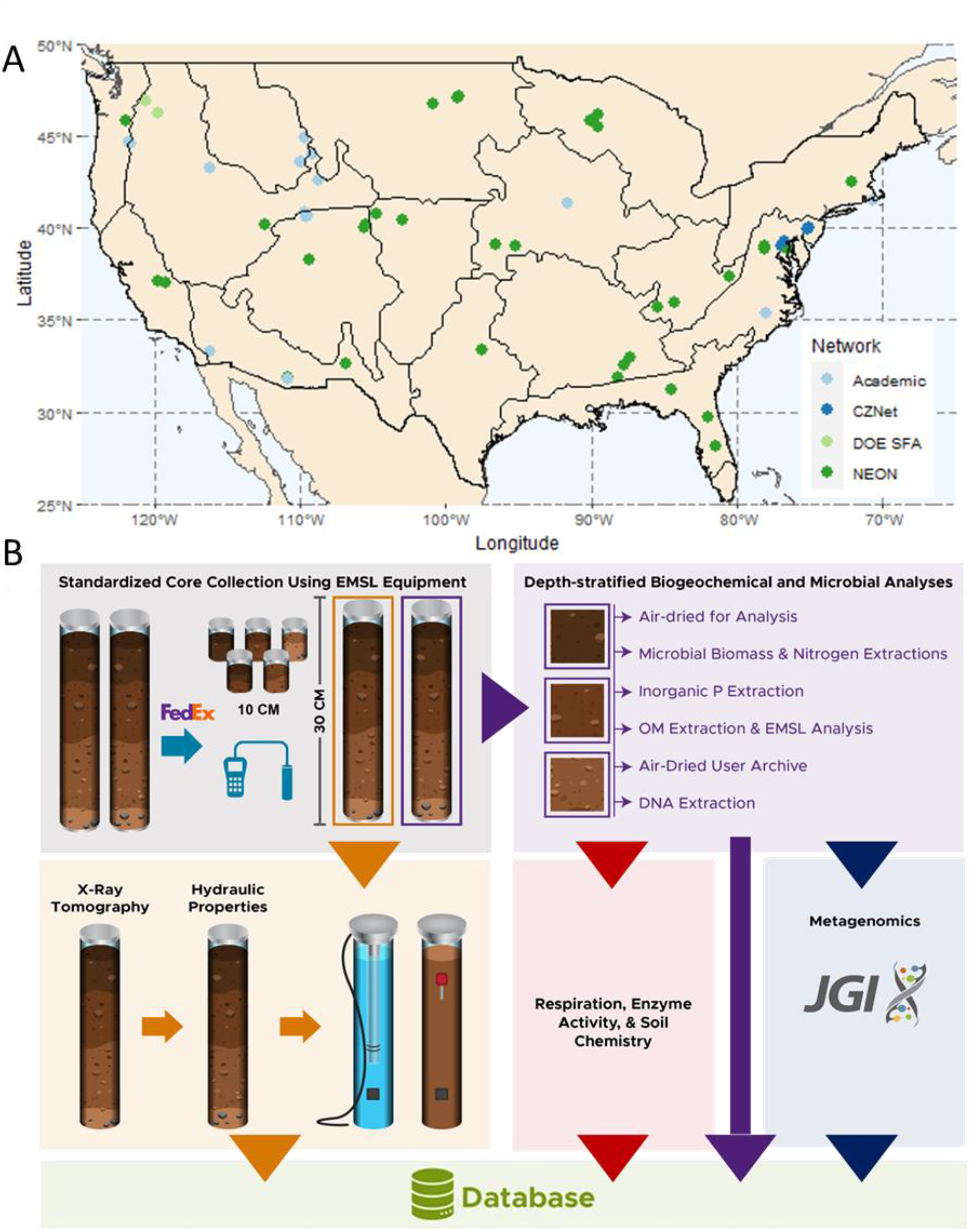
(A) Sites included in the 1000 Soils Pilot overlain on National Ecological Observatory Network (NEON) eco-domains. The color of the symbol indicates site affiliation as Academic (light blue), Critical Zone Network (CZNet; dark blue), Department of Energy Science Focus Area (DOE SFA; light green), and NEON (dark green). (B) Soil core analysis workflow.

#### Data Generation Overview

Collaborators shipped all cores to EMSL on ice within 24 hr of core collection for immediate processing. One 30-cm core was dedicated to a biotic workflow for time sensitive measurements, and the other 30-cm core was dedicated to an abiotic workflow for structural and hydraulic characterization (Figure 1B). To account for surface heterogeneity, three of the 10 cm cores were homogenized with the top 10 cm of the larger biotic core. A full list of data types generated by the 1000 Soils Pilot is available in SI Table 1.

Immediately on arrival (<48 hr), we processed the biotic core by separating it into 10 cm depth intervals, which were then sieved at 4 mm. The middle section was archived, and all analyses were conducted on homogenized top (0-10 cm) and bottom (deepest 10 cm) sections as described below. We measured microbial biomass C and N via chloroform fumigation (Brookes et al., 1985; Witt et al., 2000; Zhao et al., 2022). Aliquots of K_2_SO_4_ extractions from the initial (background) steps of microbial biomass measurements were archived at - 20° C for later analysis of NO_3_ via Cd-reduction method and NH_4_ via spectroscopic determination (Bremner, 1965; Dahnke and Johnson, 1990). Inorganic P was extracted from each soil section via Bray or Olsen extraction (depending on pH) and subsequent analysis via methods outlined in (Bray and Kurtz, 1945; Olsen et al., 1982). We collected sub-samples of soil for DNA extraction and subsequent metagenomic sequencing on the Illumina NovaSeq platform (stored at -80° C). We measured soil pH using a 1:1 soil to water ratio using a calibrated pH probe. Gravimetric water content was determined via oven drying at 60° C. Remaining soil from each section was air dried prior to the analyses below. Beta-glucosidase potential activity and respiration rates were measured using colorimetric assays and the CO_2_ burst method. Cation concentrations from 1:10 ammonium acetate extraction (K, Ca, Mg, Na) and 1:2 soil to diethylenetriaminepentaacetic acid (DPTA) extraction solution (Zn, Mg, Cu, Fe, B, SO_4_^2-^ - S) extraction were measured with Inductively coupled plasma mass spectrometry (ICP-MS), while SOM concentration was determined by total organic carbon-total nitrogen (TOC-TN) analyzer and composition from liquid chromatography mass spectrometry (LC-MS), and Fourier transform ion cyclotron mass spectrometry (FTICR-MS) analysis. Details on SOM characterization are provided in the SI.

The abiotic core was used to determine soil structure and hydraulic properties. We used X-ray Computed Tomography (XCT) to measure soil pore size, connectivity, distribution, and volume, as detailed below. In addition, soil hydraulic properties were measured using a series of instruments by The Meter Group (KSat, Hyprop, WP4C, Meter Group, Inc., USA).

Detailed methods for data collection and analysis for XCT and FTICR-MS can be found in the SI.

### Case Study: Impacts of Wildfire Regimes on Soil; Warm Springs, Oregon

#### Importance of fire on ecosystems

As the frequency of wildfires continue to increase due to anthropogenic climate change and land management practices, it is critical to understand how fire disturbances affect belowground biogeochemistry (Abatzoglou and Williams, 2016; Huffman and Madritch, 2018). Wildfires alter belowground processes that generally result in decreased structural stability, microbial respiration and biomass, and SOM concentration (Certini, 2005; Doerr et al., 2006; Fairbanks et al., 2020; González-Pérez et al., 2004; Matosziuk et al., 2020; Neary et al., 2005; Wang et al., 2012). For instance, Certini et al. (Certini et al., 2011) reported wildfire significantly decreased total soil C, which was associated with increased lignin degradation. Others have reported the selective loss of SOM oxygen containing functional groups and/or with less condensed structures in response to wildfire (González-Pérez et al., 2004). Changes in SOM concentration and composition are coincident with structural and hydrologic changes including decreased water infiltration and increased soil erosion (Verma and Jayakumar, 2012). Because belowground C storage and bioavailability are dramatically altered by wildfire, it is important to understand the nature of wildfire effects on SOM chemistry and accessibility. Below, we describe a case study to demonstrate the utility of high-resolution data for addressing this need.

#### Dataset Description

Our primary intent in this paper is to stimulate discussion and interest in large-scale collaborative efforts by introducing MONet. To facilitate this discussion, we present initial results from three of the first cores in the 1000 Soils Pilot, collected in November 2021 in partnership with the Confederate Tribes of Warm Springs in Warm Springs, OR, USA. This region typically experiences cold, snowy winters followed by warm dry summers with a mean annual precipitation of 1778 mm and a mean annual precipitation of 8.3 °C. Soil samples were collected from plots dominated by warm and moist type grand fir (*A. grandis*) and snowbrush (*C. velutinus*). The unburned (UB plots) had forb and shrub ground cover, mature living conifers, and surface litter present (>1mm) and had no recent burn history in the last 20 years. The moderate burn frequency (MB) plots contained shrubs and dead-standing trees but very little surface litter (<1 mm) and had moderate severity burns in 2014 and 2020. The high frequency burn (HB) plots consisted of a low shrub ground cover, no live or dead standing trees, and very little surface litter (<1 mm) and had moderate severity burns in 2003, 2014 and 2020.

This subset of data is intended as a proof of concept for the utility of high-resolution data in understanding impacts of fire burn histories on soil physio-chemical properties and microbial functions. Though the dataset is small and does not allow for robust statistics (n = 3), it provides a wealth of information that is unfeasible with traditional biogeochemical approaches. With these differences in mind, we compared FTICR-MS and XCT across burn histories and soil depth to show differences in molecular SOM composition and soil structure.

#### Soil Biogeochemistry and Microbial Activity

We first compared soil biogeochemical properties including, β-glucosidase activity (BG), microbial biomass C (MBC) and N (MBN), and respiration at 24 and 96 hr for differences among soil sections and burn histories (Fig 2A and 2B). Elevated microbial biomass concentrations, respiration rates, and potential enzyme activities in the unburned site are consistent with increased biological availability when compared to the burned sites (Fairbanks et al., 2020). We found the highest respiration rate at UB followed by the MB, with the lowest respiration at HB (Figure 2B). This is consistent with other literature, showing the sterilizing effects of high frequency fires on microbial respiration (Ajwa et al., 1999; Lombao et al., 2021). Across the first 24 hours after re-wet, the rate of CO_2_ produced was greater in the top 10 cm vs. the lowest 10 cm across all cores. However, after 96 hours the rate of CO_2_ production in soils from the bottom 10 cm was greater than the top 10 cm for all sites (Fairbanks et al., 2020). Similarly, β-glucosidase potential activity and microbial biomass C and N were highest at UB (SI Figure 1A and Figure 2B respectively). Lower microbial activity at the burned sites may be related to chemical recalcitrance or physical protection of SOM related to the fires (Ajwa et al., 1999; Lombao et al., 2021). For all the sites, respiration increased between 24 and 96 hours in the bottom 10 cm, possibly reflecting less bioavailable SOM in deeper soils than in surface soils (Fang and Moncrieff, 2005). Corresponding changes in soil chemistry are shown in SI Figure 2.

**Figure 2:**
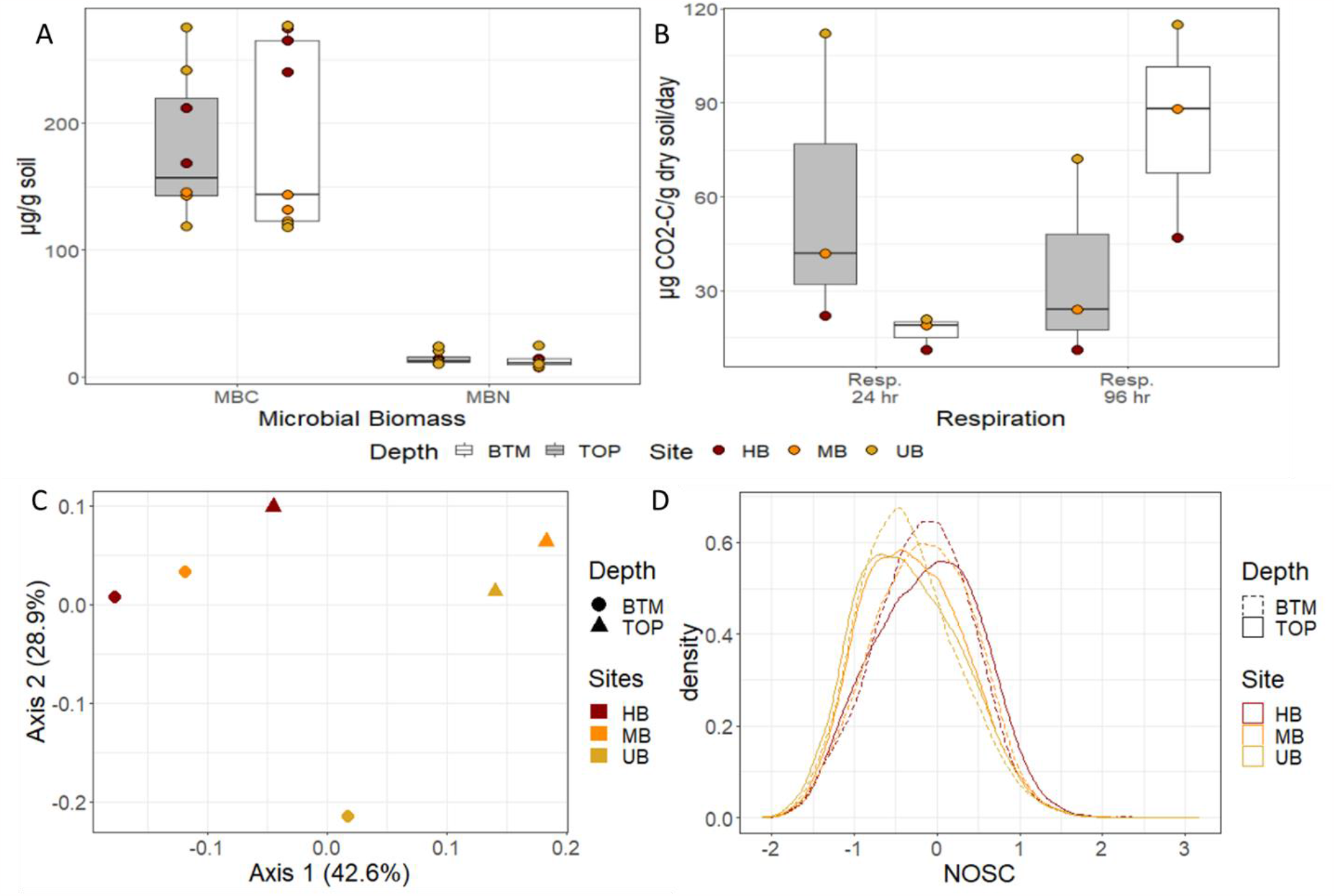
A) Microbial biomass C (MBC) and N (MBN) concentrations, B) respiration (24 and 96 hr; μg CO_2_-C/g dry soil/day), C) SOM composition, and D) NOSC of SOM profiles. A and B are boxplots where the median is represented by a line, the edges of the boxes reflect 25^th^ and 75^th^ percentile, and the end of the whiskers reflect the minimum and maximum. C shows a principal component analysis of SOM composition, and D displays a density curve of NOSC values for each sample. For all plots, UB, MB, and HB are denoted in yellow, orange, and red respectively. In A and B, data from top and bottom sections are denoted by grey and white boxes respectively. In C, section is denoted by shape, and in D, section is denoted by dashed vs. solid lines.

#### X-ray Computed Tomography

To examine pore space heterogeneity across soil sections and burn histories, we used XCT to interrogate the top and bottom 10 cm of each soil core. We compared total porosity, pore connectivity, and mean pore volume between the top and bottom sections of cores from UB, MB, and HB. Wildfire can impact soil structure at least to 10 cm depth depending on its duration, frequency, and intensity (Doerr et al., 2006). For UB and HB, we observed higher porosity and pore connectivity in the top sections when compared to the bottom sections. This is likely related to the higher abundance of root systems in the top 10 cm. However, there was minimal difference between total porosity or connectivity between the top and bottom section of the MB site, possibly due to a large root section resulting in a larger mean pore size (SI Figure 5 and SI Table 2). Across all cores, total porosity was more variable in the top than bottom section (8.82-20.08% vs 6.52-8.49%; SI Table 2). Pore connectivity was highest in the top section at UB (96.41%) and lowest in the bottom section at UB (70.12%); potentially indicating a homogenizing effect of wildfire on pore network structure at MB and HB. Overall, these data suggest that wildfire may decrease pore connectivity and total porosity in surface soils, likely contributing to increased hydrophobicity and soil erosion, while the deeper soils are more insulated from wildfire effects.

#### Soil Organic Matter Composition

Soil organic matter is derived from chemical, physical, and biological processes that decompose plant and animal residues into microbial metabolites and other byproducts (Jobbágy and Jackson, 2000; Köchy et al., 2015; Lehmann and Kleber, 2015; Schmidt et al., 2011; Tarnocai et al., 2009). This generates a heterogenous mixture of compounds in SOM pools. We used FTICR-MS to distinguish differences in SOM chemistry along depth profiles and across the three wildfire histories. There was a clear separation in water-extractable SOM composition between the top and bottom sections of each core as well as across burn regimes, as indicated by principal component analysis (Figure 2C). Additionally, the bottom sections exhibited more similar SOM pools (i.e., tighter clustering) than the top sections, possibly signifying stronger impacts of wildfire on surficial soils.

Since fire results in the oxidation of SOM to carbon dioxide (CO_2_), we hypothesized that the extent of SOM oxidation, as indicated by the nominal oxidation state of carbon (NOSC), would be related to burn regime. In the top section, the average NOSC of SOM increased with stronger burn regimes (−0.34, -0.29, and -0.11 respectively for UB, MB, and HB), indicating more oxidization, while there was a binary effect of burning on NOSC in the bottom section. MB and HB had similar mean NOSC that was higher than UB (−0.16 vs -0.35). The presence of more oxidized SOM in burned soils, is likely related to the incomplete oxidation of SOM and the addition of charred materials from wildfire (Schmidt and Noack, 2000; Velasco-Molina et al., 2016; Watson et al., 2005). While limited to three sites, changes in NOSC could be an indicator of previous fire exposure, with the NOSC becoming more oxidized with increased high frequency fire exposure that persists even into deeper soil horizons.

#### Inferred Biochemical Transformations and Microbial Metabolism of SOM

We also analyzed FTICR-MS spectra to understand potential biochemical transformations and microbial pathways involved in SOM decomposition. Commonly inferred biochemical transformations such as the addition of methyl chain (-CH_2_) groups were abundant at all sites and depths. We also note that one of the most abundant transformations observed was the conversion of carboxyl groups to CO_2_ (greater than 1400 occurrences at each site and depth). The transformation of carboxyl groups to CO_2_ likely results in combustion byproduct associated with incomplete oxidation of SOM during fire exposure. Within each core, the oxidation and/or addition of CO_2_ accounted for a greater proportion of transformations at depth when compared to the top section at the same site. This is potentially related to the translocation of water-soluble oxidized OM to deeper soil sections (Hockaday et al., 2007).

Additionally, we mapped compounds detected by FTICR-MS to microbial reference pathways in KEGG to infer mechanisms of SOM decomposition. Across all samples, nearly 50% of all mapped metabolites were detected in a set of eight metabolic pathways. These included ‘biosynthesis of secondary metabolites’ (map01110, 11.7%), ‘metabolic pathways’ (map01100, 8.4%), ‘biosynthesis of antibiotics’ (map01130, 7.5%), and ‘microbial metabolism in diverse environments’ (map01120, 4.9%) among other pathways (SI Table 4). Notably, the degradation of aromatic compounds (map012220) and the polycyclic aromatic hydrocarbons (PAH) degradation (map00624) were also among pathways with the highest number of mapped metabolites (16th and 19th most metabolites). The degradation of aromatic compounds was the most common pathway in the core with high frequency burns (HB). The degradation of PAHs was most common in the MB site; 1.61% of metabolites mapped to metabolic pathways in the moderate frequency burned core were associated with the degradation of PAHs in comparison to <1% of mapped metabolites in the unburned core.

#### Case Study Conclusions

Overall, our results were consistent with literature describing wildfire impacts on soils and provided greater understanding of molecular changes in soil organic matter composition and structure than is possible with typical measurements. In soils with recent fire history, we observed decreases in soil respiration, microbial biomass, and potential enzyme activity (Certini, 2005; Fairbanks et al., 2020; Guénon et al., 2013; Lombao et al., 2021; Neary et al., 2005; Wang et al., 2012). This was associated with shifts in SOM composition, with higher rates of oxidation apparent in surface soils.

Deeper soils also showed increased NOSC as a function of previous fire history. By inferring biogeochemical transformations from SOM pool composition, we also show the possible conversion of carboxyl groups to CO_2_ that is likely related to the oxidizing effects of wildfire. Using metabolic pathway mapping, we observed that the degradation of aromatic compounds and polycyclic aromatic hydrocarbons may also be associated with wildfire. Finally, wildfire appeared to induce structural changes including decreased pore connectivity and total porosity in surface soils, possibly contributing to hydrophobicity and decreased soil infiltration which can result in greater soil erosion (Certini, 2005; Verma and Jayakumar, 2012). The combined molecular and microstructural changes observed in this use case provide important insight and considerations in the management and modelling of forest ecosystems post-wildfire. As the MONet database continues to expand, we hope to use this information to model wildfire impacts on belowground C cycling at the regional to CONUS scale.

### Towards a CONUS-scale high resolution molecular SOM database

Earth system models currently rely on soil attributes such as soil texture, moisture, and C concentration to parameterize C fluxes. This highly simplified approach results in high levels of uncertainty at the pore to core scale. Molecular information has the potential to reduce this uncertainty through rapidly improving multiscale models (Blankinship et al., 2018; Feng et al., 2021; Kramer and Chadwick, 2018; Nave et al., 2021; Quesada et al., 2020; Rasmussen et al., 2018; Viscarra Rossel et al., 2019; Von Fromm et al., 2021; Yu et al., 2021).

To achieve these goals, MONet is developing a database of high-resolution soil organic matter, metagenomic, and structural information from soil cores across the CONUS. We collaborate with existing databases and networks, as well as with individual researchers, using standardized sample collection and high throughput processing pipelines, metadata documentation, and raw and processed open data publication. The focus of our data generation is on variables that are key to the parameterization of hierarchical multiscale models beginning with microbially explicit models and culminating in ESMs.

In this article, we present the 1000 Soils Pilot for MONet including a use case to highlight the utility of high-resolution measurements for understanding the impacts of wildfires on soils. While the statistical analysis is limited by the small sample number (n=3), we observed trends in soil biogeochemical values, soil structure, and SOM composition related to fire history and soil section that are consistent with existing literature. We are currently accepting requests for collaboration, and we anticipate that all data from the first set of 76 cores will be available in 2023. With the support and collaboration of the scientific community, we aim to facilitate the next generation of fundamental knowledge and model representations of soil C cycling, which in turn constrain uncertainties in the global climate.

## Acknowledgments

Collection of soils from the Warm Springs Reservation was funded by a Washington State University, Vancouver Mini-Grant to SSP and KM, and a Washington State University College of Arts and Sciences Seed Grant to SSP, KM, TC, and EG. In addition, we would like to thank the tribal leaders from the Confederate Tribes of Warm Springs. This research was performed on a project award (https://dx.doi.org/10.46936/intm.proj.2021.60141/60000423) from the Environmental Molecular Sciences Laboratory, a DOE Office of Science User Facility sponsored by the Biological and Environmental Research program under Contract No. DE-AC05-76RL01830. This material is based upon work supported by the U.S. Department of Energy (DOE), Office of Science (SC), Biological and Environmental Research (BER), Research Development and Partnership Pilot (RDPP), under Award Number(s) DE-SC0023150.

This report was prepared as an account of work sponsored by an agency of the United States Government. Neither the United States Government nor any agency thereof, nor any of their employees, makes any warranty, express or implied, or assumes any legal liability or responsibility for the accuracy, completeness, or usefulness of any information, apparatus, product, or process disclosed, or represents that its use would not infringe privately owned rights. Reference herein to any specific commercial product, process, or service by trade name, trademark, manufacturer, or otherwise does not necessarily constitute or imply its endorsement, recommendation, or favoring by the United States Government or any agency thereof. The views and opinions of authors expressed herein do not necessarily state or reflect those of the United States Government or any agency thereof.

