## Supplemental Information for "One thousand soils for molecular understanding of belowground carbon cycling"

**Affiliation:**

**Standardized sampling protocol:**

1000 Soils Pilot Field Sampling Protocol

Written by Maggie Bowman

This protocol describes soil core sampling for the 1000 Soils Pilot Campaign. Please keep in mind that it is the collector’s responsibility to ensure compliance with any environmental regulations associated with sampling (e.g. permits and site access). After QA/QC, data generated by the 1000 Soils Project will be publicly available according to FAIR principles (<https://www.go-fair.org/fair-principles/>), and we encourage data collectors to abide by CARE principles surrounding indigenous peoples (<https://www.gida-global.org/care>). Upon receipt, please verify that this kit contains the materials listed below. Sampling should be completed between Thursday and Saturday and shipped overnight to EMSL between Sunday and Tuesday the following week using the provided return shipping labels. Please contact Maggie Bowman with any questions.

**Summary of Field Work and Samples Collected**

As part of the 1000 Soils field sampling, you will be collecting meter measurements (temperature, volumetric water content, and electrical conductivity), soil samples (3” x 12” Abiotic/Biotic paired cores and 2” x 4” smaller cores), and infiltration measurements.

**Identify Sampling Location**

Identify one sampling location that is as representative of the landscape as possible. When possible, samples should be collected with ~100 m of an eddy covariance tower or climate monitoring station. Please note a short description of the sampling location about the surrounding landscape on the metadata sheet (e.g., upslope, downslope, North facing, etc.).

**Prior to Sample Collection**

Before collecting samples, please ensure that the kit contains all materials listed below. In addition, please place the blue ice provided for shipping in the freezer and fill the two included bottles with 444 ml of water each for infiltration measurements.

**Materials**

| **1000 Soils sampling kit should contain the following:** | **Items provided by you:** |
| --- | --- |
| 1. Insulated shipping container 2. Freezer blue ices packs (please place in freezer for minimum 48 hours prior to return shipping) 3. Return shipping labels 4. Nitrile gloves (bags containing small, medium, and large gloves) 5. Clipboard containing hard copy of data sheet, writing utensils, and tape 6. 3”x 12” Coring tubes x4 and Caps x8 7. Four 3” core catchers (if needed) 8. 2” x 4” Coring tubes (4 per site) 9. AMS slide hammer 10. AMS Intact Corer 11. AMS Soil Recovery Auger Corer (SRA) 12. AMS Extender (2’) 13. AMS Auger Handle 14. Wretch 15. Wooden block 16. Hammer/Mallet 17. Pool Noodle (used to stabilize cores/fill open space in coring tubes during shipping) 18. 2 bags per site core to prevent leaking 19. Tape measure 20. Box cutter 21. Ear protection 22. Tape (black electrical for sealing tubes and orange electrical for labelling if needed) 23. Meter group PROCHECK + Teros 12 meter 24. Zip top plastic storage bags 25. Infiltration Ring 26. Plastic Wrap 27. Two 500 ml Nalgene Bottle | 1. Cooler and blue ice (separate form shipping blue ice) to keep samples cold in the field 2. Method for collecting latitude and longitude in decimal degrees in the field (this can be done using a smart phone) 3. Method for taking pictures of field site and sample cores (this can be done using a smart phone) 4. Stopwatch (can use phone app) 5. 500 ml of water 6. -20 C freezer to freeze ice packs prior to shipping (do not freeze cores) 7. Refrigerator to store cores prior to shipping 8. Access to a FedEx shipping location to return packages. 9. Packing tape to close shipping kit. 10. Optional field measurements (please include measurements, methods and units on the data sheet if applicable). |

**Collect Metadata**

1. Record the sample kit number, general vegetation type (e.g., conifer forest or tall prairie), general weather conditions (sunny, rainy, extreme heat, etc.), longitude and latitude in decimal degrees (smart phone app ‘My GPS Coordinates’), and the time and date of sampling.
2. It the kit is received damaged, but useable, please make a note of this on the metadata sheet.
3. Take pictures of the sampling location prior to sampling. Please include pictures of the soil with the measuring tape extended to 1 meter and 3 additional pictures of the ground cover, overlying vegetation, and landscape.
4. Please ensure that all metadata is collected prior to leaving the field.

**Collect Impact Soil Cores**

1. Within the selected area, identify 2 specific locations within 1 meter of each other for replicate cores, and put on the provided nitrile gloves.
2. Brush away any surface litter and debris using the stainless-steel bench scraper to cut a square into the surface soil and remove the vegetation layer. Do not remove the organic soil horizon. Place the removed litter layer in the provided Ziploc bag.
3. Locate one of the large core liners, remove the end caps and pool noodle (if inside coring tubes and store these items in a clean dry place). With the empty core tube place, it vertically into the impact coring barrel.
   a. Abiotic cores will be used to measure soil pore structures and hydraulic properties, it is imperative that the soil core remain as intact as possible. To do this we will be using a pool noodle to prevent the core from separating after collection.
4. Place the coring tube into the impact coring barrel with the catch cup at the tip of the corer. Screw the corer onto the slide hammer using the 2’ extension if needed.
   a.
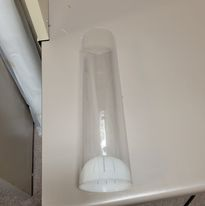
Depending on soil texture, you may need a core catcher (white half sphere). For soils that fall out of the coring tube, remove the liner, and insert the coring cup into the base of the liner as pictured.
   b. **To prevent damage to the equipment, ensure that all connections are secure and do not force them to screw together this can result in cross threading the equipment.**
5. Place the tip of the coring barrel on the ground and hammer the soil corer into the ground by sliding the hammer up and striking down.
6. Hammer the soil corer until no further progress is made until no further progress can be made (. Remove the slide hammer and attach the handle to turn the core and pull the core up.
   a. If you are still having trouble removing the core, rock the core left and right to loosen the core from the surrounding soil and remove the core vertically by pulling upward.
   b. If you feel the soil will not stay in the core liner during removal even with the soil catcher installed, dig adjacent to core, and insert a trowel horizontally at the bottom of the core prior to removal from the ground.
7. Remove the cap at the top of the soil corer and pull the core liner out maintaining the vertical orientation. Cap both the top and bottom of the core liner with the caps provided.
   a. If having issues removing the core liner from the coring device, you can push up gently on the edges of the core liner from the
8. Trim the pool noodle to fill the remaining space in the coring tube using the provided box cutter. Cap the top of the tube and secure both the top and bottom caps with the provided black electrical tape.
9. Label the tube with the site code and “Core A”. Please denote which end is the top of the soil core.
10. Repeat steps 4-7 with the tube labelled “Core B” (coring tube without pool noodle, biotic core). Capping the top of Core B without filling the gap.
11. Keeping cores vertical, take pictures of both cores with the measuring tape and record the total depth of each core on the metadata sheet.

**Collecting Auger Cores**

1. If the above coring methods do not work for your soils, you can collect cores using the soil recovery auger (SRA).
2. Place a soil core liner with the soil catcher in the bottom into the SRA and cap the auger with the open attachment.
3. Attach the SRA to the 2’ extender and handle.
4. Using the provided mallet, hammer the corer into the soil about 2 cm to start the corer.
5. Once started turn the auger handle clockwise to collect a core. Stop once the auger can no longer core or the auger has reached the full depth.
6. Follow the above instructions to remove the auger and core.

**Collecting Surface Cores**

1. To collect surface cores using the smaller 2” diameter cores remove the litter layer from a spot within 1 m of Core A and Core B. Label the 4 mini cores Core C1, C2, C3, and C4.
2. Place the coring liner into the soil and using the hammer and block of wood to hammer in the coring liner.
3. Rock the core back and forth to loosen and start to release it and pull the core liner and core up. If the core can not be easily removed use the trowel to dig the core out.
   a. It is helpful to cap the top of the core while still in the soil. This acts as a vacuum as well as a point of leverage.
4. Cap the mini cores and tape the ends with electrical tape.
5. Place the sealed mini cores in a bag.

**Contingency Plans and Alternative Methods**

1. If the above coring methods do not work for your soils, you can collect cores using any coring methods that work at your field site. Please indicate on the metadata sheet that Impact soil cores failed, and in the notes, section provide details on the coring alternative. If possible, maintain the approximate vertical structure of the core (e.g., mineral vs. organic layers). Note whether approximate spatial structuring was maintained on the metadata sheet.
2. If no other coring methods exist, an alternative method for collecting soils is to use a trowel or shovel to dig a soil pit and pack a core with the material removed doing your best to maintain the soil depths Like above, please indicate on the metadata sheet that Impact soil cores failed, and in the notes, section provide details on the coring alternative. If possible, maintain the approximate vertical structure of the core (e.g., mineral vs. organic layers). Note whether approximate spatial structuring was maintained on the metadata sheet.

**Collect Field Measurements**

1. At the center point between the two soil cores, collect measurements for temperature, moisture and electrical conductivity.
2. Start by removing the Meter Group Procheck and Teros 12 from the case.
3. Insert the Teros 12 serial cable into the top port on the Procheck meter.
4. Using the Menu button, turn on the Procheck and press Enter to start measuring.
5. Remove the foam cover over the 3 metal probes (be careful the probes are sharp)
6. On the Measurement tab ensure that the instrument is set up for mineral soils (this wetting will be used for all soil types and the word mineral will appear on the bottom of the screen).
7. Gently press the probes into the ground, if there is any resistance do not force the probes. Move the probe over a few inches and try again.
8. Once the probe is in the ground, allow the readings to stabilize for 3 minutes.
9. Save the data by pressing the “SAVE” button and using the up and down arrows to select characters and numbers using the enter button to advance to the next space. Save the data using the 4 character site code followed by a “-“ and 0001 or 0002.
   a. This will help keep multiple measurements from the same site/project separate.
10. Press Save one final time to store the data on the ProCheck meter.
    a. For example, PROS-0001 and PROS-0002 for 2 measurements at the Prossor site.
11. Record the save name and measurements on the metadata sheet.

**Infiltration Measurements**

1. Clear the sample area of debris and place the 6” ring on the soils. Using the block of wood and hammer drive the 6” ring into the soil. Drive the ring into the ground about 1”.
2. Within the soil use your finger to firm up the soil around the inside edges of the soil ring.
3. Line the top of the ring with the provided plastic wrap to prevent disturbance to the soil when adding the water.
4. Add 444 ml of water from the plastic bottles. Poor the water into the ring lined with plastic wrap.
5. Remove the wrap leaving just the water in the ring and start recording the time using a stopwatch. Record the amount of time (in minutes) it takes for the water to infiltrate the soil. Stop timing when the soil surface is just glistening.
6. In the same ring, repeat steps 2-4 for a second infiltration measurement.
   a. If infiltration measurements take >30 minutes for the initial round do not completed a second round of water additions and not on the metadata sheets provided that

<https://www.nrcs.usda.gov/wps/PA_NRCSConsumption/download?cid=nrcseprd1358428&ext=pdf>

**Transport and Store Samples After Collection**

1. After cores are collected, transport them on blue ice to a refrigerator (4 °C) and store vertically when possible.

**Upload Metadata and Photos**

1. Send images of the metadata sheets and soil cores to with the header 1000 Soils – Site – Metadata in the subject line.

**Shipping**

1. All cores should be stored in the refrigerator until they are shipped on a Monday or Tuesday. It is critical that overnight shipments be made no later than a Wednesday. Please don’t ship later in the week in case there are shipping delays. We cannot receive cores over the weekend. Contact before you sample/ship if this is not possible. It is vital to store samples vertically in the refrigerator or ship within 72 hours following sampling, though 48 hr is preferable. It can be useful to pack the cooler as close as is reasonable to the time FedEx will ship the package. This maximizes the time cores stay cold.
2. When you’re ready to ship, place the cores in the provided large zip top bags to prevent leaking.
3. Pack all remaining materials and the soil probes as received, wrapping all items in the provided packing materials (See photos attached). Place the frozen “blue ice” on top.
4. Place the insolated lid on top and attach the metadata sheet in a plastic bag to the outside of the Styrofoam lid and seal the box to keep it dry.
5. Please don’t return used gloves or other waste generated. Please dispose of the waste appropriately.
6. Tape the outer box closed. Please use enough packing tape to make sure it won’t open during shipping.
7. Adhere the provided shipping label to the outside of the box by removing the backing from the plastic sleeve that contains the return shipping label.
8. Drop the package off at FedEx, or have it picked up by FedEx (must ensure same day pick up).
9. On the same day you ship the package, notify that you shipped the package, and include the FedEx tracking number in your email. This is critical to ensure sample integrity and timely delivery. The subject line of the email should be “1000 SOILS SHIPPED SAMPLES – [Sample kit ID #].

**Check List**

- **Record the sample kit number on the metadata sheet**
- **Collect site specific metadata**
  - **Date**
  - **Time**
  - **GPS**
  - **Site full name**
  - **Individuals present**
  - **Vegetation**
  - **Weather**
- **Pictures of the sampling location**
- **Collect surface litter**
- **Collect Core A (abiotic impact core)**
- **Collect Core B (biotic impact or auger core)**
- **Collect Cores C1-4 (mini cores collected at the surface)**
- **Collect and store meter measurements**
  - **Temperature**
  - **Moisture**
  - **Electrical Conductivity**
- **Infiltration time**
  - **1^st^ infiltration**
  - **2^nd^ infiltration**

**Metadata Sheet**

**Site Metadata**

Sample Kit ID: ______________________________________________________

Site Name: _________________________________________________________

Date (YYYYMMDD): ________________ Time (HH:MM AM/PM):______________

GPS Method: _______________________________________________________

Lat: ____________________________ Long: _____________________________

Cores Collected by: __________________________________________________

__________________________________________________________________

Vegetation: _________________________________________________________

Weather: __________________________________________________________

**Meter Measurements**

Save Name: _______________________ Temperature: _____________________

Save Name: _______________________ Vol. Water Content: ________________

Save Name: _______________________ Bulk Density: ______________________

**Coring Notes**

Core A Notes: _______________________________________________________

__________________________________________________________________

___________________________________________________________________

Core B Notes: _______________________________________________________

__________________________________________________________________

___________________________________________________________________

Core C1-3 Notes: ____________________________________________________

__________________________________________________________________

___________________________________________________________________

**Infiltration Measurements**

Infiltration Time 1 (MM:SS): _______________

Infiltration Time 2 (MM:SS): _______________

Notes: ____________________________________________________________

**Additional Notes**

___________________________________________________________________

___________________________________________________________________

___________________________________________________________________

___________________________________________________________________

___________________________________________________________________

___________________________________________________________________

___________________________________________________________________

___________________________________________________________________

___________________________________________________________________

___________________________________________________________________

___________________________________________________________________

___________________________________________________________________

___________________________________________________________________

**SI Table 1:** List of measurements included in the Molecular Observation Network sampling.

| **Measurement** | **Method** | **Analysis** |
| --- | --- | --- |
| Metagenomics | Illumina NovaSeq | In house extractions; External sequencing partners |
| Texture | Hydrometer | External Partners |
| Respiration | CO_2_ Burst, 24 and 96 hrs. | External Partners |
| Potential Enzyme Activity  (β-glucosidase) | Colorimetric Assay | External Partners |
| Ortho-P | Bray/Olsen Method | In house extractions; External Partners |
| NO_3_^-^ and NH_4_ | 0.5 M K_2_SO_4_ Extract; Colorimetric | In house extractions; External Partners |
| Geochemistry – K, SO_4_-S, B, Zn, Mn, Cu, Fe, Ca, Mg, Na; Total Bases; Cation Exchange Capacity | Ca, Mg, Na, K – S-5.1 (Ammonium Acetate Method) | External Partners |
|  | Zn, Mn, Cu, Fe, B, SO_4_-S – S6.11 (DPTA Extraction) S-10.10 (Ammonium Replacement Method) |  |
| Total Carbon and Nitrogen | AOAC 972.3 AOAC 990.3 | External Partners |
| Gravimetric Water Content (GWC) | 60 C for 48 hours or until no change in mass | EMSL |
| Microbial Biomass C and N | Chloroform fumigation | EMSL |
| pH | 1:1 soil water ratio | EMSL |
| Water Extractable Organic Matter (WEOM) – TOC/TN and Composition | Shimadzu TOC/TN analyzer LC-MS (Q Exactive 2) FT-ICR-MS (Solid Phase Extraction; Scimax 2xR 7T) | EMSL |
| Soil Pore Network Structure | X-ray Computed Tomography Low resolution scan of full core (~70 um) High resolution scans of soil layers (~30 µm) | EMSL |
| Soil Hydraulic Properties | KSAT, HYPROP2, and WP4C (The Meter Group) | EMSL+A6:C20A2:C20A1:C20BA8:C20 |

**SI Methods:**

*Experimental Methodology for Soil Physical Characterization by X-ray Computed Tomography.*

The intact abiotic soil core was imaged using XCT on an X-Tek/Metris XTH 320/225 kV scanner (Nikon Metrology, Belmont, CA, USA). Data was collected at 110 kV and 325 μA X-ray power. A 0.5-mm thick Cu filter was used to enhance image contrast by blocking out low-energy X-rays. The core was rotated continuously during the scans with momentary stops to collect each projection (shuttling mode) while minimizing ring artifacts. A total of 2000 projections were collected over 360˚ rotation recording 2 frames per projection with 500 ms exposure time per frame. Image voxel size was 70.1 microns. The whole core was imaged in two segments/tiles to increase spatial resolution.

Higher resolution images were collected by zooming in on both the top and bottom sections 10 cm of the core. These data were also collected at 110 kV and 325 μA X-ray power with a 0.5-mm thick Cu filter. A total of 3142 projections were collected over 360˚ rotation recording 2 frames per projection with 708 ms exposure time per frame. Image voxel size was 33.6 microns.

*Data Analysis for Soil Physical Characterization by X-ray Computed Tomography.*

Soil pore size and connectivity were calculated from XCT images. The images were reconstructed to obtain three-dimensional datasets using CT Pro 3D (Metris XT 2.2, Nikon Metrology). Representative slice and 3D images were created using VG Studio MAX 2.1 (Volume Graphics GmbH, Heidelberg Germany) and Avizo 2019.2 (Thermo Fisher Scientific, Waltham, MA). Image processing and porosity analysis was carried out using Avizo 2019.2. The reconstructed 3D volume data was filtered using a median filter and cropped to exclude artificially created pore space that occurred during sampling. The final volume data processed for porosity segmentation was 1300 x 1200 x 1500 voxels to exclude surrounding air space while maximizing the analyzed soil volume.

Each image was segmented into pore or water space vs. soil based on manual thresholding. Each voxel was assigned a value between 1 and 2 that correlates to its assigned component (i.e., pore/water or soil).

Determining pore connectivity requires the selection of parameters that a pore network must adhere to in order to be considered “connected” and part of the main pore network. Pore voxels were used to determine pore network connectivity. Using the Axis Connectivity Module in Avizo, we required pore networks to extend between two opposing faces of the volume’s bounding box, and two planes (X, Y, or Z axis) were chosen depending on which method is most suitable to the core sample. In cases where no volumes satisfied the axis connectivity requirements, the connected pore network was defined as the connected group of pores with the largest volume using the Filter by Measure module in Avizo. Distinguishing individual pores and pore throats required the use of the watershed algorithm, separating a connected region into distinct objects (Sarkar and Siddiqua, 2016).

Following segmentation, the following parameters were extracted (1) pore volumes from connected and unconnected pores, (2) pore coordination numbers, and (3) pore throat diameters. These data were obtained for soil macropores down to 70 microns for whole core imaging and 38.1 microns for higher resolution imaging.

*Experimental Methodology for Soil Organic Matter Characterization by FTICR-MS.*

SOM was extracted from soils using 6 g of air-dried soil with 30 mL of ultrapure DI water. Soil water mixtures were shaken for 2 hours at 800 rpm. After shaking, soil mixtures were centrifuged at 6000 rpm for 8 minutes. The supernatant was collected and divided into aliquots for TOC/TN, LC-MS and FTICR-MS. Aliquots collected for TOC/TN and LC-MS were filtered using an ultrapure DI rinsed 0.45 um PES filter and stored frozen until analysis. TOC/TN was determined via Shimadzu TOC-TN analyzer. The aliquot for LC-MS was lyophilized and concentrated to 50x using an 80:20 methanol:water mixture prior to analysis. LC-MS spectra were collected. Concentrated OM samples were analyzed using the Q-Exactive 2 with, using a Hilic column, samples and run in both positive and negative mode. FTICR-MS samples were acidified to pH 2 using concentrated phosphoric acid and salts were removed using Agilent Bond Elut PPL SPE cartridges.

A 7 Tesla (7T) Bruker ScimaX Fourier transform ion cyclotron resonance mass spectrometer (FTICR-MS) (Bruker, SolariX, Billerica, MA, USA), was used to collect high-resolution mass spectra of the SOM water extracts. Solutions were injected directly into the instrument using a custom automated direct infusion cart that performed two offline blanks between each sample. The FTICR-MS was outfitted with a standard electrospray ionization (ESI) source, and data was acquired in negative mode with the needle voltage set to +4.0kV. Data is collected from 150 m/z – 1000 m/z at 8M. three hundred scans were co-added for each sample and internally calibrated using OM homologous series separated by 14 Da (–CH_2_ groups). The mass measurement accuracy was typically within 1 ppm for singly charged ions across a broad m/z range (150 m/z - 1100 m/z). Bruker Data Analysis (version 5.0) was used to convert raw spectra to a list of m/z values by applying FTMS peak picker module with a signal-to-noise ratio (S/N) threshold set to 7 and absolute intensity threshold to the default value of 100. Chemical formulae were then assigned using Formularity (Tolić et al., 2017), an in-house software, following the Compound Identiﬁcation Algorithm (Kujawinski and Behn, 2006; Minor et al., 2012; Tfaily et al., 2017). Chemical formulae were assigned based on the following criteria: S/N >7, and mass measurement error < 0.5 ppm, taking into consideration the presence of C, H, O, N, S and P and excluding other elements. Data analysis procedures for FTICR-MS are described in the SI.

*Data Analysis for Soil Organic Matter Characterization by FTICR-MS.*

Biochemical transformations and microbial metabolic pathways potential of SOM decomposition were inferred from FTICR-MS spectra, following procedures previously described by (Danczak et al., 2020; Garayburu-Caruso et al., 2020a; Garayburu-Caruso et al., 2020b; Graham et al., 2018; Graham et al., 2017; Kaling et al., 2018; Moritz et al., 2017) amongst others. Inferred biochemical transformations were based on a protocol initially published by (Breitling et al., 2006). This approach is feasible with FTICR-MS data because the set of peaks in each sample are related by measurable and clearly defined mass differences corresponding to gains and losses of compounds. Briefly, all possible pairwise mass differences were calculated between every peak in a sample and compared to a reference list of 1255 masses associated with common biochemical transformations (within 1 ppm). Differences were inferred to represent the gain or loss of a compound with the specified mass. For example, a mass difference of 99.07 corresponds to a gain or loss of the amino acid valine, while a difference of 179.06 corresponds to the gain or loss of a glucose molecule.

In parallel, possible microbial metabolic pathways in each sample were identified by locating chemical formulae assigned to *m*/*z*'s within metabolic pathways defined in the Kyoto Encyclopedia of Genes and Genomes (KEGG, [http://www.kegg.jp](http://www.kegg.jp/)) (Graham et al., 2018; Kanehisa and Goto, 2000; Tfaily et al., 2018). Chemical formulae were mapped to KEGG pathways using R code available at https://github.com/EMSL-Computing/ftmsRanalysis to detect all KEGG pathways containing a given formula. For example, a peak with an assigned formula C20H16O9 and mapped to KEGG pathway “map00254” (Aflatoxin biosynthesis) which contains C20H16O9 as an intermediate. However, only a subset of compounds detected by FTICR-MS are defined within the KEGG database, because peaks must be assigned a chemical formula and that chemical formula must be present in a KEGG pathway.

Finally, because wildfire can result in incomplete combustion and oxidation of SOM (González-Pérez et al., 2004), we calculated the nominal oxidation state of C (NOSC) to represent the energy required to oxidize different molecules of organic matter, calculated as $NOSC= -(\frac{4C+H-3N-2O+5P-2S}{C})+4$(Koch and Dittmar, 2006; 2016; LaRowe and Van Cappellen, 2011). NOSC is calculated from the assigned chemical formula and can range from -4 (reduced) to +4 (oxidized), with individual compounds in organic matter typically ranging from -3 to +2 (Keiluweit et al., 2016).

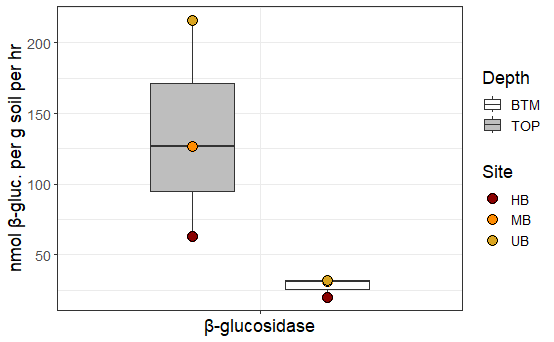

**SI Figure 1:** β-glucosidase (β-gluc; nmol β-glucosidase per g soil per hr) concentration for top (grey) and bottom (BTM; white) sections of soil cores for unburned (UB; yellow), moderate burn (MB; orange), and high burn (HB; red) frequency. The edges of the boxes reflect 25^th^ and 75^th^ percentile, and the end of the whiskers reflect the minimum and maximum.

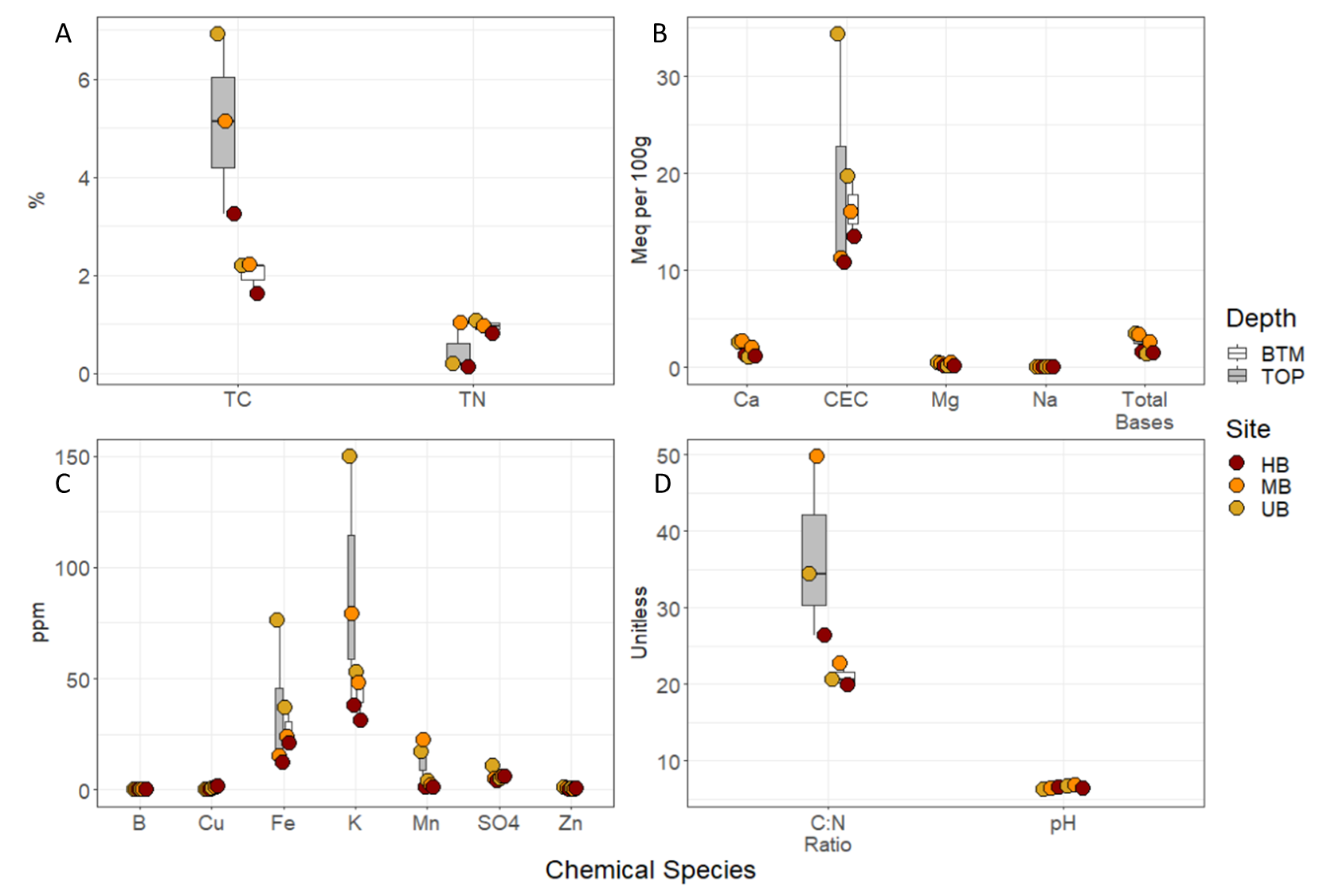

**SI Figure 2:** A) Percent total carbon (TC) and total nitrogen (TN) concentrations, B) Calcium (Ca), Magnesium (Mg), Sodium (Na), cation exchange capacity (CEC), and total bases concentrations (Meq per 100 grams of soil), C) Boron (B), copper (Cu), Iron (Fe), potassium (K), manganese (Mn) sulfate (SO_4_-S), and zinc (Zn) concentration in ppm D) C:N ratio and pH, the edges of the boxes reflect 25^th^ and 75^th^ percentile, and the end of the whiskers reflect the minimum and maximum. For all plots, UB, MB, and HB are denoted in yellow, orange, and red respectively. Additionally top and bottom sections are denoted by grey and white boxes respectively.

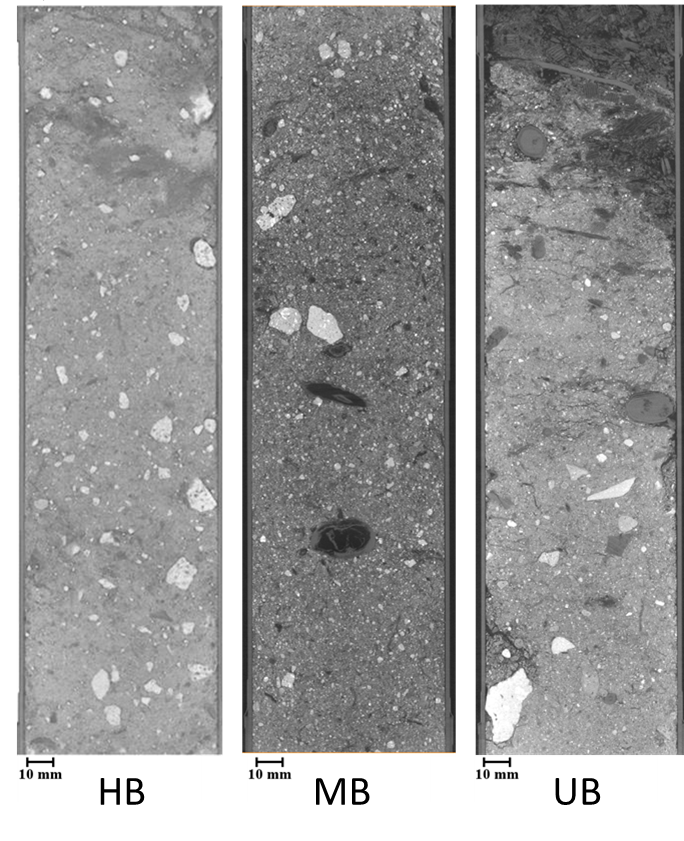

**SI Figure 3:** Low resolution full core XCT images for high burn (HB) frequency, moderate burn (MB) frequency, and unburned soils.

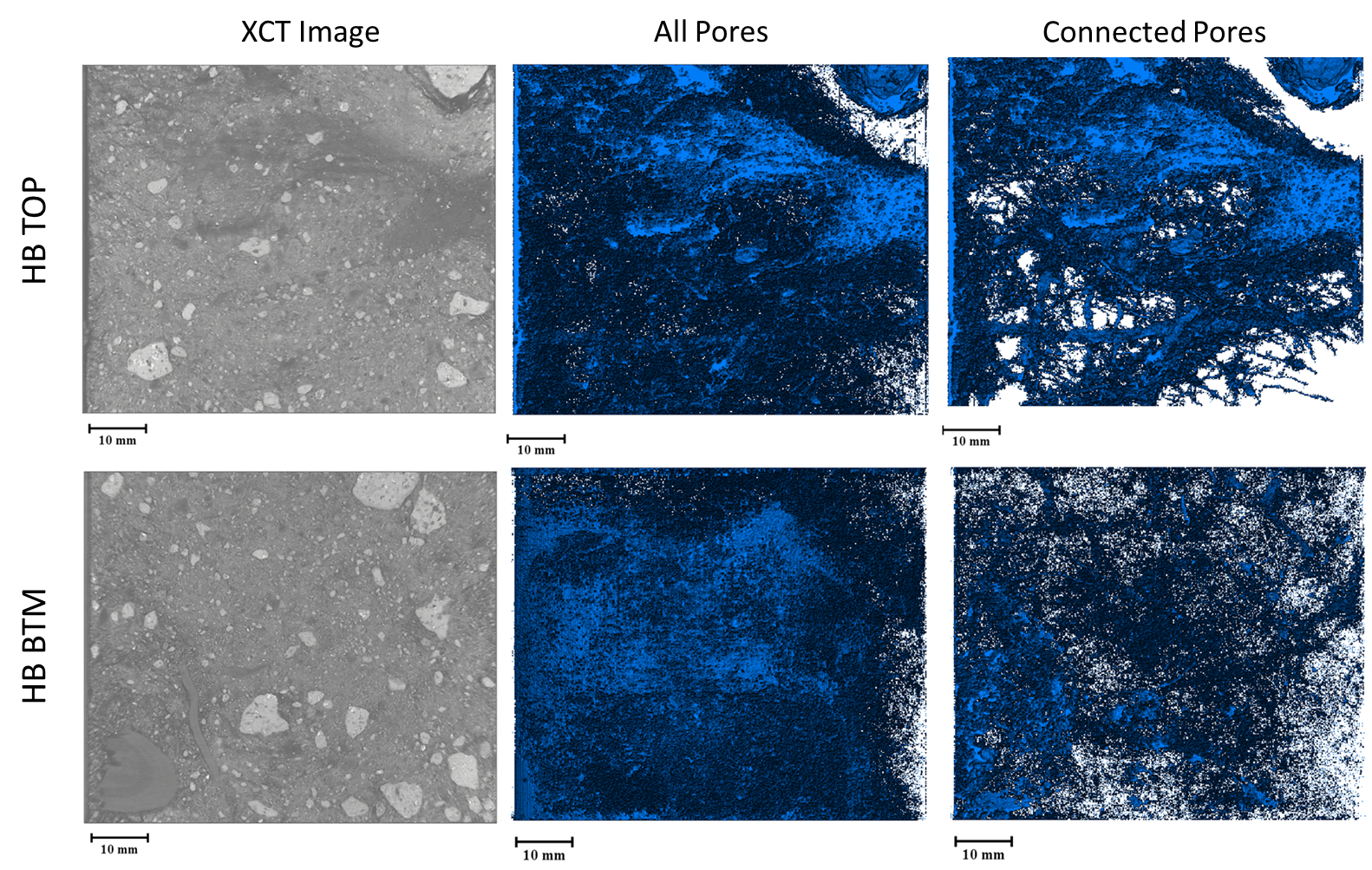

**SI Figure 4:** High resolution XCT images (left column), all detected pores (center column), and connected pores (right column) for top and bottom sections of the high burn (HB) frequency soil core.

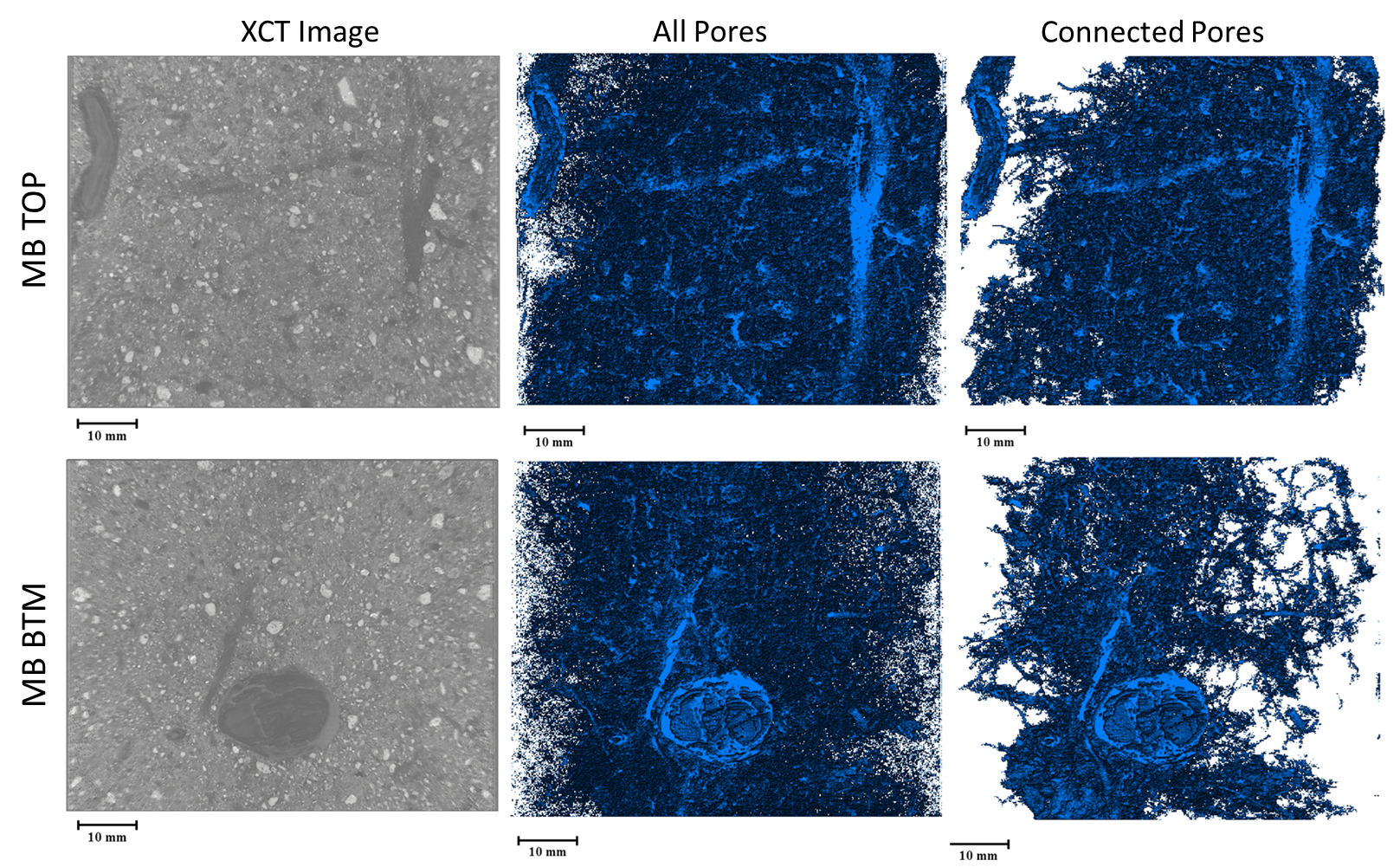

**SI Figure 5:** High resolution XCT images (left column), all detected pores (center column), and connected pores (right column) for top and bottom sections of the moderate burn (MB) frequency soil core.

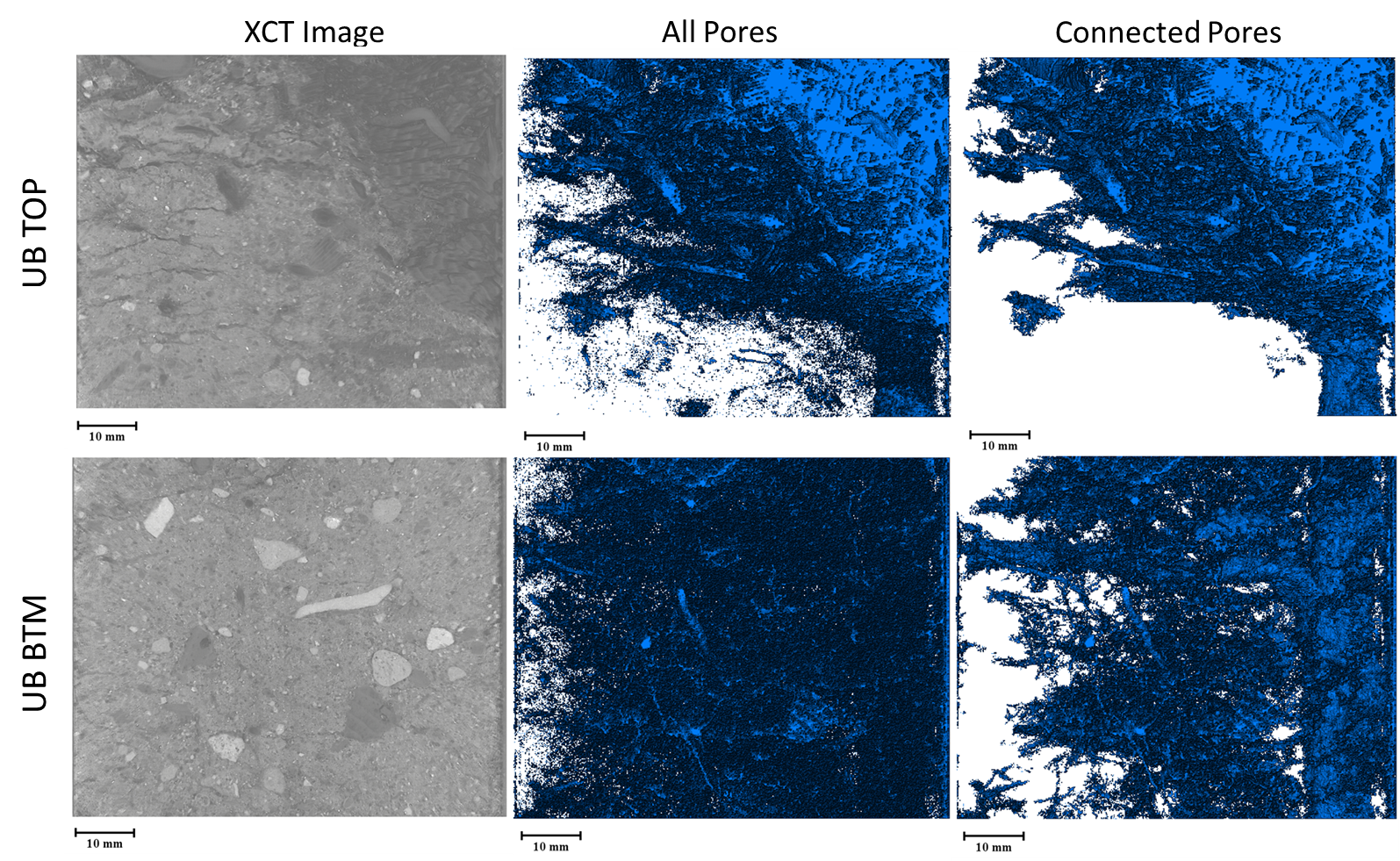

**SI Figure 6:** High resolution XCT images (left column), all detected pores (center column), and connected pores (right column) for top and bottom sections of the unburned (UB) soil core.

**SI Table 2:** Pore characteristics calculated from cropped (50x48x60 mm) high resolution X-ray computed tomography (XCT) images for top (TOP) and bottom (BTM) sections for unburned (UB), moderate burn (MB) frequency, and high burn (HB) frequency soil cores.

| Site | Depth | Connected Pores | Pore Size | | | | | Pore Volume | | Wet Bulk Density |
| --- | --- | --- | --- | --- | --- | --- | --- | --- | --- | --- |
|  |  |  | Min | Max | Mean | Median | Variance | Mean | Fraction |  |
|  |  | % | mm | mm | mm | mm | mm | mm^3^ | % | g/mm^3^ |
| HB | TOP | 90.41 | 0.047 | 32.153 | 0.072 | 0.060 | 0.002 | 0.007 | 14.11 | 1.89 |
| HB | BTM | 83.84 | 0.047 | 26.636 | 0.077 | 0.060 | 0.003 | 0.006 | 7.75 | 1.78 |
| MB | TOP | 82.49 | 0.047 | 26.821 | 0.079 | 0.060 | 0.003 | 0.005 | 8.82 | 1.63 |
| MB | BTM | 80.09 | 0.047 | 26.578 | 0.081 | 0.063 | 0.004 | 0.007 | 8.49 | 1.00 |
| UB | TOP | 96.41 | 0.047 | 37.022 | 0.073 | 0.060 | 0.003 | 0.024 | 20.08 | 1.63 |
| UB | BTM | 70.12 | 0.047 | 22.794 | 0.076 | 0.060 | 0.003 | 0.003 | 6.52 | 1.90 |

**SI Table 3:** Pore size distribution count from cropped (50x48x60 mm) high resolution X-ray computed tomography (XCT) images for top (TOP) and bottom (BTM) sections for unburned (UB), moderate burn (MB) frequency, and high burn (HB) frequency soil cores.

| Site | Depth | Pore Size Distribution | | | | | | | | | | | |
| --- | --- | --- | --- | --- | --- | --- | --- | --- | --- | --- | --- | --- | --- |
|  |  | 0-0.1 | 0.1-0.2 | 0.2-0.3 | 0.3-0.4 | 0.4-0.5 | 0.5-0.75 | 0.75-1.0 | 1.0-5.0 | 5.0-10.0 | 10.0-25.0 | 25.0-50.0 | >50.0 |
|  |  | mm | mm | mm | mm | mm | mm | mm | mm | mm | mm | mm | mm |
| HB | TOP | 886705 | 138367 | 17010 | 4145 | 1303 | 861 | 141 | 41 | 0 | 0 | 1 | 0 |
| HB | BTM | 868024 | 150471 | 21597 | 5323 | 1742 | 1095 | 216 | 107 | 0 | 0 | 1 | 0 |
| MB | TOP | 844866 | 171447 | 23709 | 5485 | 1703 | 1097 | 185 | 81 | 0 | 0 | 1 | 0 |
| MB | BTM | 833063 | 178802 | 26197 | 6460 | 2201 | 1380 | 310 | 160 | 0 | 0 | 1 | 0 |
| UB | TOP | 899380 | 126857 | 15815 | 4011 | 1346 | 895 | 173 | 96 | 0 | 0 | 1 | 0 |
| UB | BTM | 856422 | 160567 | 22995 | 5459 | 1773 | 1091 | 173 | 92 | 1 | 1 | 0 | 0 |

**SI Table 4:** SOM decomposition inferred from mapped compounds detected by FTICR-MS to microbial reference pathways in KEGG.

*Table uploaded separately*
